# Maternal circulating syncytiotrophoblast-derived extracellular vesicles contain biologically active 5’-tRNA halves

**DOI:** 10.1101/721795

**Authors:** William R Cooke, Adam Cribbs, Wei Zhang, Neva Kandzija, Carolina Motta-Mejia, Eszter Dombi, Rannya Ri, Ana Sofia Cerdeira, Christopher Redman, Manu Vatish

## Abstract

The placenta releases syncytiotrophoblast-derived extracellular vesicles (STB-EV) into the maternal circulation throughout gestation. STB-EV dependent signalling is believed to contribute to the widespread maternal adaptive physiological changes seen in pregnancy. Transfer RNA (tRNA) halves have been identified in vesicles released from other human and murine organ systems, which alter gene expression in target cells. Here, we characterise tRNA-half expression in STB-EV and demonstrate biological activity of a highly abundant tRNA-half. Short RNA from ex-vivo, dual-lobe placental perfusion STB-EV was sequenced, showing that most (>95%) comprised tRNA species. Whole placental tissue contained <50% tRNA species, suggesting selective packaging and export of tRNA into STB-EV. Most tRNA within STB-EV were 5’-tRNA halves cleaved at 30-32 nucleotides. The pattern of tRNA expression differed depending on the size/origin of the STB-EV; this was confirmed by qPCR. Protein synthesis was suppressed in human fibroblasts when they were cultured with a 5’-tRNA half identified from STB-EV sequencing. This study is the first to evaluate tRNA species in STB-EV. The presence of biologically active 5’-tRNA halves, specific to a vesicular origin, suggests a novel mechanism for maternal-fetal signalling in normal pregnancy.

## Introduction

During normal pregnancy, placental syncytiotrophoblast, the epithelium coating the chorionic villi, releases extracellular vesicles into the maternal circulation [1]. The syncytiotrophoblast of a normal placenta has a surface area of ≈ 10 m^2^ and releases more than 3 g of syncytiotrophoblast-derived extracellular vesicles (STB-EV) each day at term [2]. Levels of STB-EV in the peripheral venous blood of pregnant women increase with gestational age and fall sharply post-partum [2,3]. STB-EV can be classified by size, mode of release and contents. Small extracellular vesicles (SEV), also known as exosomes, are around 50-200 nm diameter and are synthesised intracellularly and released into the maternal circulation from multivesicular bodies [1]. Medium-large vesicles (MLEV), also known as microvesicles, are around 200-1000 nm diameter and are produced by blebbing from the cell surface. STB-EV contribute to a spectrum of immunological, cardiovascular and endocrine changes which occur in adaptation to normal pregnancy [4].

MicroRNAs within STB-EV are differentially expressed between normal and diseased pregnancies, suggesting a possible signalling role [5]. *In vitro* microRNAs can be delivered into the mitochondria and endoplasmic reticulum of endothelial cells [6]. In culture, trophoblast vesicles attenuate viral replication in target cells via microRNAs [7]. Work on microRNAs dominate the field; however one study has reported transfer RNA (tRNA) species in STB-EV isolated by placental explant culture: 60-65% small RNA sequences were miRNAs, 20-22% were ribosomal RNA (rRNA) and 13-17% were tRNAs [5]. This latter finding was not further investigated.

tRNAs transfer amino acids to ribosomes during translation. They are 70-90 nucleotides long and have a “cloverleaf” secondary structure (Figure 1D). tRNA breakdown products were identified in the urine of patients with malignancies suggesting potential use as biomarkers [8]. Subsequently, a variety of post-transcriptional modifications in tRNAs were identified comprising two distinct groups of tRNA species: tRNA halves and tRNA fragments (tRFs) [9]. tRNA halves are produced by the ribonucleases angiogenin and RNase T [10]. Two molecules of around 31-40 nucleotides are produced when the anticodon loop is cleaved: those from the 5’ end of the whole tRNA are named 5’-tRNA-halves, while 3’-tRNA-halves are derived from the 3’ end. tRFs are shorter species produced by a variety of nucleases which cleave tRNAs at several nucleotides to produce fragments 15-25 nucleotides long [11]. They are also named tRF-5 for those derived from the 5’ end of the mature tRNA, tRF-3 for those from the 3’ end and tRF-1 for those from tRNA precursor molecules.

**Figure 1.**
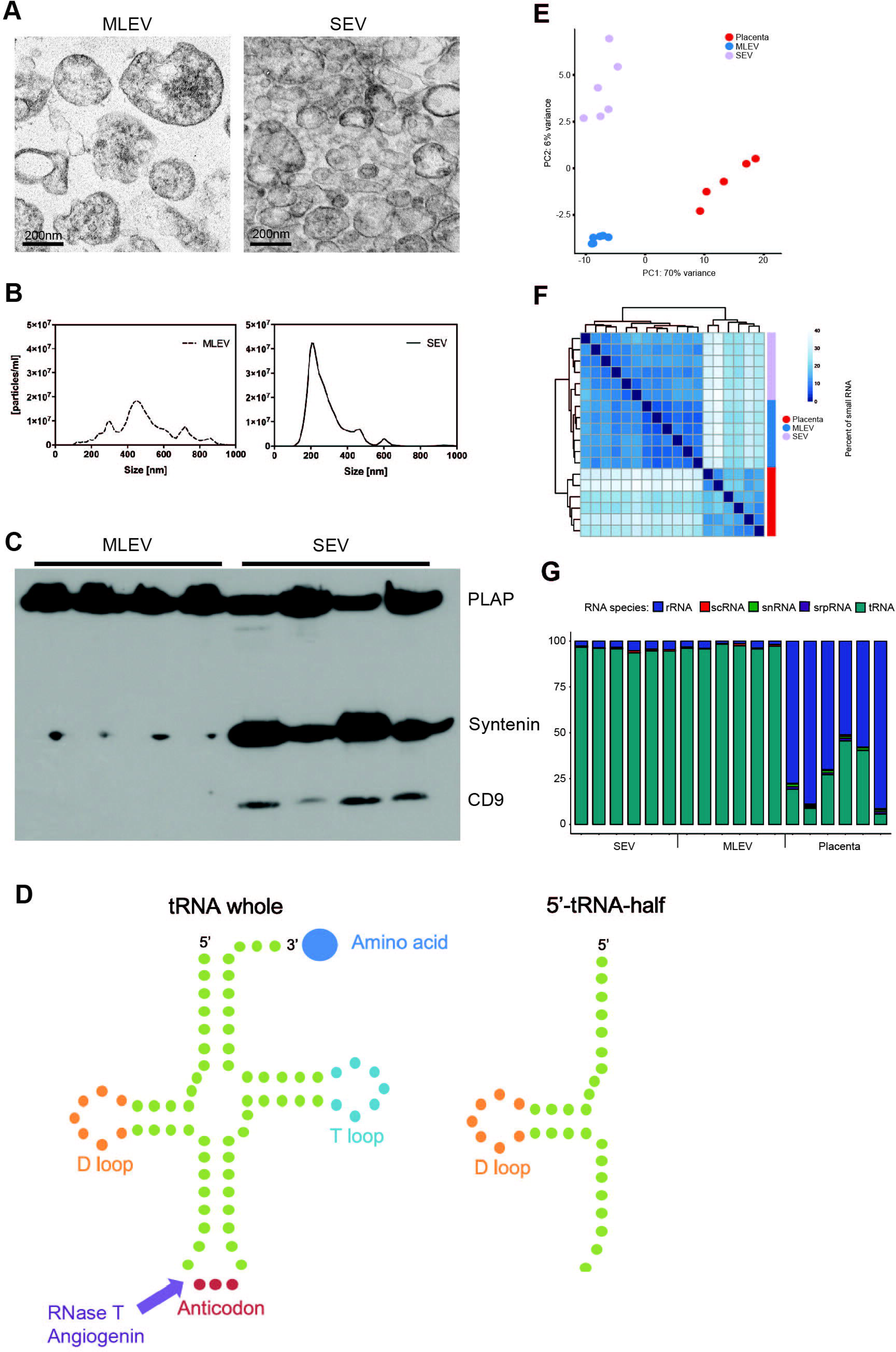
A: Transmission electron microscope (TEM) images showing medium large extracellular vesicles (MLEV; left) and small extracellular vesicles (SEV; right). B: nanoparticle tracking analysis (NTA) profile for medium large extracellular vesicles (MLEV; left) and small extracellular vesicles (SEV; right). C: Representative immunoblot of medium large extracellular vesicles (MLEV; left) and small extracellular vesicles (SEV; right) showing expression of placental alkaline phosphatase (PLAP; 60 kDa band) in all samples; Syntenin (32 kDa band) and CD9 (25 kDa band) in SEV samples only. D: Schematic diagram showing the secondary “clover leaf” structure of a whole tRNA and demonstrating a point of cleavage to create a 5’-tRNA-half. E: Principal component analysis demonstrating clear separation of RNA source tissue by tRNA species. F: Sample-sample distances are displayed as a heatmap. The heatmap was generated by DESeq2 software package and shows the Euclidean distance between each sample. G: Distribution of sequencing reads in SEV, MLEV and whole placental libraries mapped to all RNA species.

tRNA halves have been identified in different tissues and bodily fluids, including seminal fluid, sperm, serum and T cells [12–15]. They are signalling molecules, altering gene expression when injected into cells or inhibited with antisense oligonucleotides [13]. They can alter cell phenotype by interfering both with translation and transcription. tRNA-halves displace translation initiation factor 4E (eiF4E) resulting in translational arrest [10,16]. tRFs are also postulated to act as miRNAs: they are associated with argonaute proteins in a similar fashion as miRNAs, and have seed sequences complementary to RNA targets allowing transcriptional repression [17].

We aimed to quantify the diversity and relative expression of tRNA species in STB-EV and sought evidence that the tRNA species most abundant in STB-EV are biologically active.

## Materials and methods

### Human subjects

The project was approved by the Central Oxfordshire Research Ethics Committee (07/H0607/74 and 07/H0606/148). Participants provided written informed consent. Six placentae were collected at the time of elective caesarean section (for breech presentation or previous caesarean section) and perfused immediately. Prior to perfusion, a biopsy of placental tissue was taken from a non-perfused lobe for downstream RNA processing. Participants had healthy singleton pregnancies at >39 weeks gestation.

### Isolation and characterisation of STB-EV

Placental perfusate from a dual placental lobe perfusion system was centrifuged to pellet fractions containing medium-large extracellular vesicles (MLEV) (10,000 × g) and small extracellular vesicles (SEV) (150,000 × g) as previously described [18] for all six placentae. The MLEV and SEV were divided into aliquots and those not immediately used were frozen at −80°C. Fresh aliquots of MLEV and SEV were fixed with 4% glutaraldehyde in 0.1M phosphate buffer and processed for routine transmission electron microscopy using a Jeol 1200EX electron microscope (JEOL, Massachusetts, USA). Samples were post-fixed in 2% osmium tetroxide in 0.1M phosphate buffer; dehydrated in ethanol; treated with propylene oxide; embedded in Spurr’s epoxy resin; cut and then stained with uranyl acetate and lead citrate. A NanoSight NS500 system (Malvern Instruments, Malvern, UK) was used to measure the size and concentration of further aliquots of STB-EV using Nanoparticle Tracking Analysis (NTA). Western blotting was performed using 6 μg STB-EV. Samples were treated with HEPES lysis buffer 1:1 (50mM HEPES, pH 7.5, 2% SDS, 10% glycerol). Western blots were incubated overnight with mouse monoclonal anti-PLAP (placental alkaline phosphatase) antibody (1:1000; NDOG2, in-house antibody), rabbit monoclonal anti-Syntenin antibody (ab133267, Abcam, Cambridge, UK) and rabbit monoclonal anti-CD9 antibody (ab92726, Abcam, Cambridge, UK). Blots were developed using HRP-conjugated anti-mouse IgG antibody (1:4000) and Pierce ECL Western Blotting Substrate (Thermo Fisher Scientific, Paisley, UK). The protein concentration of isolated STB-EV was determined by BCA protein assay and STB-EV were stored at −80°C until use.

### RNA isolation

RNA was isolated from STB-EV and placental tissue using Total RNA Purification Plus Kit (Norgen Biotek Corporation, ON, Canada) following manufacturer’s instructions. RNA size and integrity were confirmed using RNA 6000 Pico kit for the Agilent 2100 Bioanalyzer (Agilent Technologies, Waldbronn, Germany) and Nanodrop™ ND-1000 spectrophotometer (Thermo Fisher Scientific, Paisley, UK).

### Next generation sequencing and library preparation

Total RNA was isolated from each sample and quantified using Nanodrop™, normalised and then libraries were prepared. Briefly, 250ng of total RNA underwent small RNA library preparation using the NEBNext Multiplex Small RNA library preparation kit (NED, Ipswich, MA, USA) according to the manufacturer’s instructions. The libraries were then checked for size and concentration using a High Sensitivity D1000 ScreenTape (Agilent Technologies, Lanarkshire, UK) to enable precise loading of the flow cell. Samples were then sequenced using an Illumina HiSeq 4000 machine.

### qPCR arrays for tRNA halves and tRNA-derived fragments

rtStar™ First-Strand cDNA Synthesis Kits (Arraystar, Maryland, USA) were used according to manufacturer’s instructions to ligate 3’ and 5’ adaptors to the end of tRNA halves and synthesise cDNA. Equal quantities of cDNA from each patient were loaded onto nrStar™ Human tRF&tiRNA PCR Array (Arraystar, Maryland, USA) to determine expression of 185 specific tRNA halves and fragments (compiled from databases and reported in recent publications). Raw CT (cycle threshold) values were normalised to the housekeeping small nuclear RNA RNU6 and compared to a reference tRNA half present in all samples (Gly-GCC-1) to determine fold tRNA expression; only tRNA species detectable with the kit’s recommended CT threshold of 35 are shown.

### Synthesised oligonucleotides

A 31 nucleotide RNA molecule corresponding to the most abundant 5’-tRNA half species identified from sequencing and PCR arrays was synthesised: 5’-tRNA-half-*Gly-GCC* (5’-GCAUUGGUGG**<u>UUC</u>**AGUGGUA**<u>GAA</u>**UUCUCGCC-3’). A scrambled RNA sequence of identical length was synthesised to act as a control, changing five nucleotides corresponding to the hairpin regions of the 5’-tRNA half in order to disrupt its secondary structure and function: *5’-tRNA-half-scramble* (5’-GCAUUGGUGG**<u>GGG</u>**AGUGGUAG**<u>GG</u>**UUCUCGCC-3’). Oligonucleotides were custom synthesised from Thermo Fisher Scientific (Paisley, UK).

### Cell culture and protein synthesis assay

Primary human fibroblasts established from skin biopsies were cultured onto optical 96-well plates at a density of 5000 cells/ml. Cells were allowed to recover overnight before medium was changed to methionine-free RPMI with an added methionine analogue (L-homopropargylglycine), using reagents from the Click-iT^®^ HPG Alexa Fluor^®^ Protein Synthesis Assay Kit (Thermo Fisher Scientific, Paisley, UK). Cells were treated with cycloheximide (40ng/ml) as a positive assay control, 500nM 5’-tRNA-half-Gly-GCC and 500nM 5’-tRNA-half-scramble. After 22 hours incubation, cells were fixed with paraformaldehyde, permeabilised with Triton-X-100 and L-homopropargylglycine molecules were detected using Alexa Fluor^®^ 488 azide. Cells were imaged using the IN Cell 1000 analyzer (GE Healthcare) using 15 fields of view per well. Images were analysed using a specialized Developer toolbox protocol to quantify mean fluorescence per cell.

### Small RNA sequencing data analysis

Raw sequencing reads were quality controlled using FastQC [19] and Fastq Screen [20] analysis and subjected to adaptor removal by Trimmomatic [21]. All analyses were performed using our tRNAnalysis pipeline (https://github.com/Acribbs/tRNAnalysis). Briefly, all reads passing filter were then subjected to mapping using the human hg38 genome build by using Bowtie1[23] (release 1.2.2) in-v1 alignment mode and best alignment stratum reporting. Initial annotation was performed against all ensemble annotated “small RNAs”, “ribosomal RNA” and other RNA annotations. We used a similar mapping strategy to Hoffman et al. [23] to specifically map the tRNA in which the tRNA precursors sequences were appended as extra ‘artificial’ chromosomes. In a first pass mapping, reads that overlap boundaries of mature tRNAs were extracted. In a second pass, the remaining reads were mapped to a tRNA-masked target genome to remove mapping artefacts.

Given that each tRNA cluster constitutes its own artificial chromosome, we used samtools [24] idxstats to count the number of reads for each tRNA cluster and merge across all samples to form one counts table. This counts table was then used for differential expression analysis applying DESeq2 [25]. Unless otherwise stated, all downstream bioinformatic analyses were performed using R 3.4 and Bioconductor. The differences between the datasets were first measured (PCA analysis) using DESeq2 (version 1.22.1). Heat maps were generated from tRNA gene-expression values using the *pheatmap* package v1.0.8 with the “ward” clustering method as default. Differences in tRNA gene expression between samples were evaluated by the Wald significance test. tRNAs with a Banjamini-Hochberg-adjusted p value of < 0.05 were considered to be significant.

### Statistical analysis

Protein synthesis assay data were analysed using GraphPad Prism 8 (GraphPad Software, California, USA). Normality of data was confirmed using D’Agostino and Pearson normality test; hypotheses were tested using an unpaired Student t-test, and values were expressed as mean ± SEM.

## Results

### Placental perfusion produces MLEV and SEV

Sequential centrifugation of eluates from the maternal side of the *ex-vivo* placental lobe dual perfusion system yielded extracellular vesicles with size profiles corresponding to MLEV (200-800 nm diameter) and SEV (150-400 nm diameter) (Figures 1A, 1B). Syncytiotrophoblast origin was confirmed by Western blotting (Figure 1C) for the presence of placental alkaline phosphatase (PLAP), a unique marker of syncytiotrophoblast. Additionally, common exosome markers (Syntenin-1 and CD9) were present only on SEV, demonstrating that this population included syncytiotrophoblast exosomes. We isolated MLEV and SEV from six healthy placentae at term. We also obtained a sample of placental tissue from a non-perfused lobe at the beginning of the perfusion from each placenta.

### More than 95% of RNA within STB-EV are tRNA species

Next, we used small RNA Next Generation Sequencing (NGS) of whole placental tissue alongside MLEV and SEV obtained by dual-lobe perfusion from the same six normal placentae. SEV and MLEV were clearly distinguished by principal component analysis (PCA) (Figure 1E) as they were in a heatmap of the sample-sample Euclidean distance (Figure 1F). Next, we quantified the number of reads mapping to different RNA species. In the placental tissue a substantial proportion of reads mapped to both rRNA and tRNA, while in both the STB-EVs, in contrast, >95% of reads mapped to tRNA (Figure 1G). MLEV and SEV contained a spectrum of different tRNAs. The most abundant were GlyGCC, GluTTC, LysCTT, GluCTC, GlyCCC, CysGCA and HisGTG (Figure 2A). Given redundancy in codons, these tRNAs can also be grouped by the amino acid they carry, of which the most abundant included glycine, glutamic acid, lysine, cysteine and histidine (Figure 2B). Whole placental tissue contained a broader variety of tRNA species, consistent with their cellular function of amino acid shuttling in translation (where all 20 amino acids will be needed).

**Figure 2.**
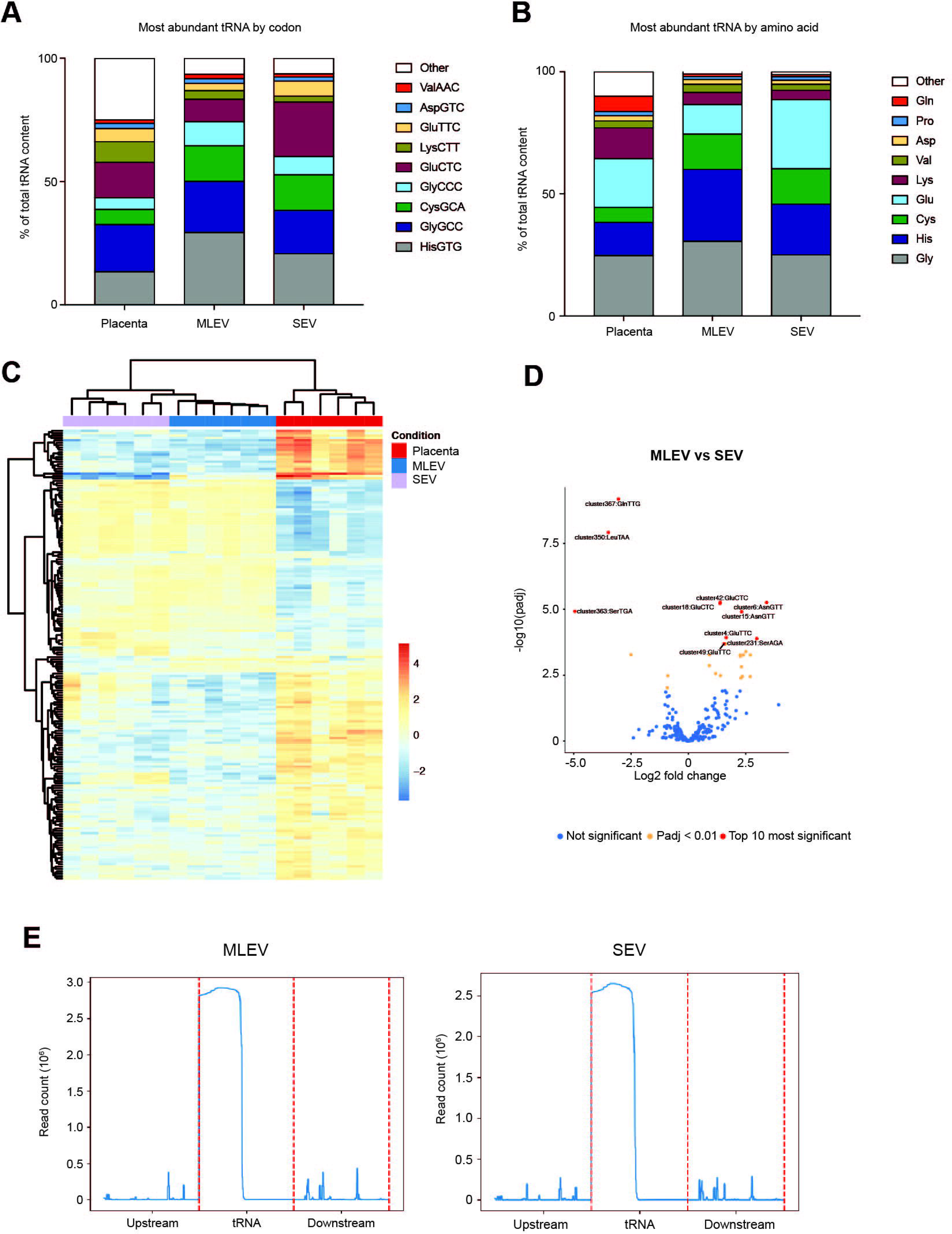
A: Proportion of sequencing reads in placenta, MLEV and SEV mapped to each tRNA and collapsed by codon. B: Proportion of sequencing reads mapped to different tRNAs within placenta, MLEV and SEV and collapsed by amino acid. C: Heatmap showing differential tRNA species expression by source tissue: placenta, MLEV and SEV. D: Volcano plot showing relative up-and down-regulation of tRNA species in MLEV when compared to SEV. E: Coverage plots showing read counts of tRNA sequences from 5’ (upstream) to 3’ (downstream) in MLEV (left) and SEV (right). The total tRNA sequence is represented between the dashed red lines. “Upstream” and “Downstream” represent other RNA sequences upstream and downstream of the tRNA.

### The tRNA profiles in placenta, MLEV and SEV showed substantial variation

The tRNA differences between placenta, MLEV and SEV were analysed by differential expression analysis, revealing 189 differential tRNA species between both types of STB-EV and placental tissue (Figure 2C). The most pronounced differences were between all STB-EV and whole placental tissue tRNA species. The patterns of tRNA species expression associated with SEV and MLEV were similar, but not identical. Despite their apparent similarities, differential expression analysis revealed 26 differentially regulated tRNAs (Figure 2D), consistent with the differences between the relative abundances of tRNA species seen in figures 2A and 2B.

### Most of the STB-EV tRNA are 5’-halves, but not 3’-halves

We next analysed coverage plots over all tRNA genes and demonstrated that most reads in both MLEV and SEV align to the 5’ end of the tRNA species, with a major cleavage point at around 30-32 nucleotides (Figure 2E).

### qPCR corroborates the presence of 5’-tRNA halves within STB-EV

In order to validate the presence of tRNA halves within the SEV and MLEV samples, we used the rtStar™ First-Strand cDNA Synthesis Kit to synthesise cDNA from tRNA halves and fragments. SEV and MLEV samples from 3 normal placentae were interrogated using a qPCR array plate containing primers for 185 recently identified tRNA halves/fragments to corroborate the NGS findings. 35/185 and 33/185 possible tRNA species were detected in MLEV samples and SEV samples, respectively (Figure 3A, 3B). The most abundant tRNA species were detected in MLEV and SEV in all samples; these corresponded to eleven 5’-tRNA-halves, one tRF-5, and one 3’-tRNA half. The most abundant 5’-tRNA halves carried the same anti-codons as those identified using RNAseq (Figure 2A): GlyGCC GluTTC, LysCTT, GluCTC, and GlyCCC. In addition to common tRNA species, there were unique findings: 19 tRNA species were only present in samples from one placenta (Figures 3A, 3B).

**Figure 3.**
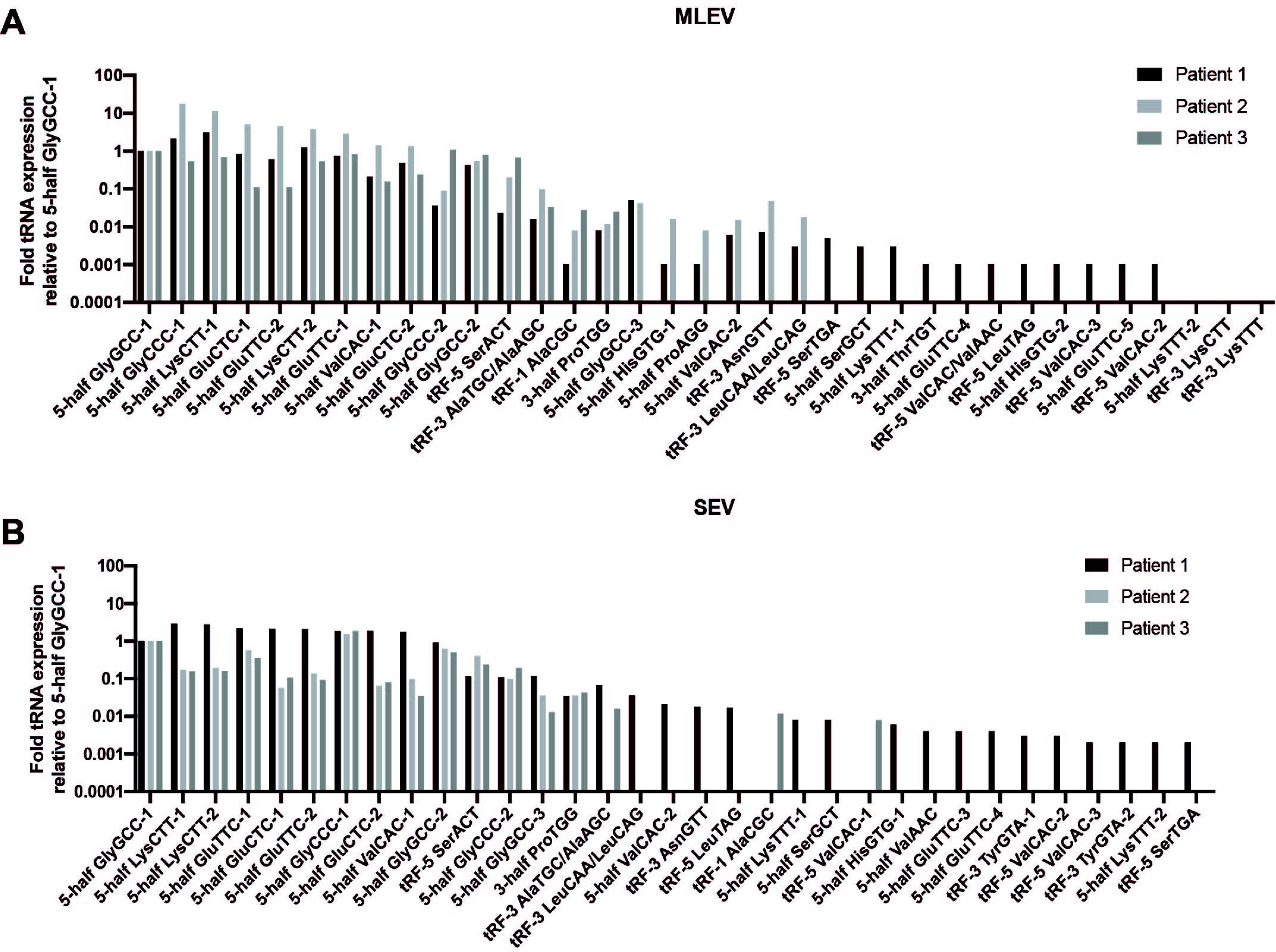
Fold tRNA half and tRNA fragment expression in medium large extracellular vesicles (MLEV; 3A) and small extracellular vesicles (SEV; 3B) relative to 5’-tRNA half Gly-GCC-1 in placental perfusion samples from three patients. Data are normalised to the housekeeping small nuclear RNA RNU6.

### Fibroblasts demonstrate biological activity of 5’-tRNA-halves

Given the variety of tRNA species identified within STBEV, we elected to investigate the *in vitro* effects of one of the most abundant (5’-tRNA-half-GlyGCC) using fibroblasts in cell culture as a model. Cells treated with 500 nM 5’-tRNA-half-GlyGCC demonstrated a mean 12.4% reduction in global protein synthesis compared to those treated with 500 nM 5’-tRNA-*half-scramble* (Figure 4). There was no significant difference between global protein synthesis in untreated (blank) cells and those treated with 500 nM 5’-tRNA-half-scramble.

**Figure 4.**
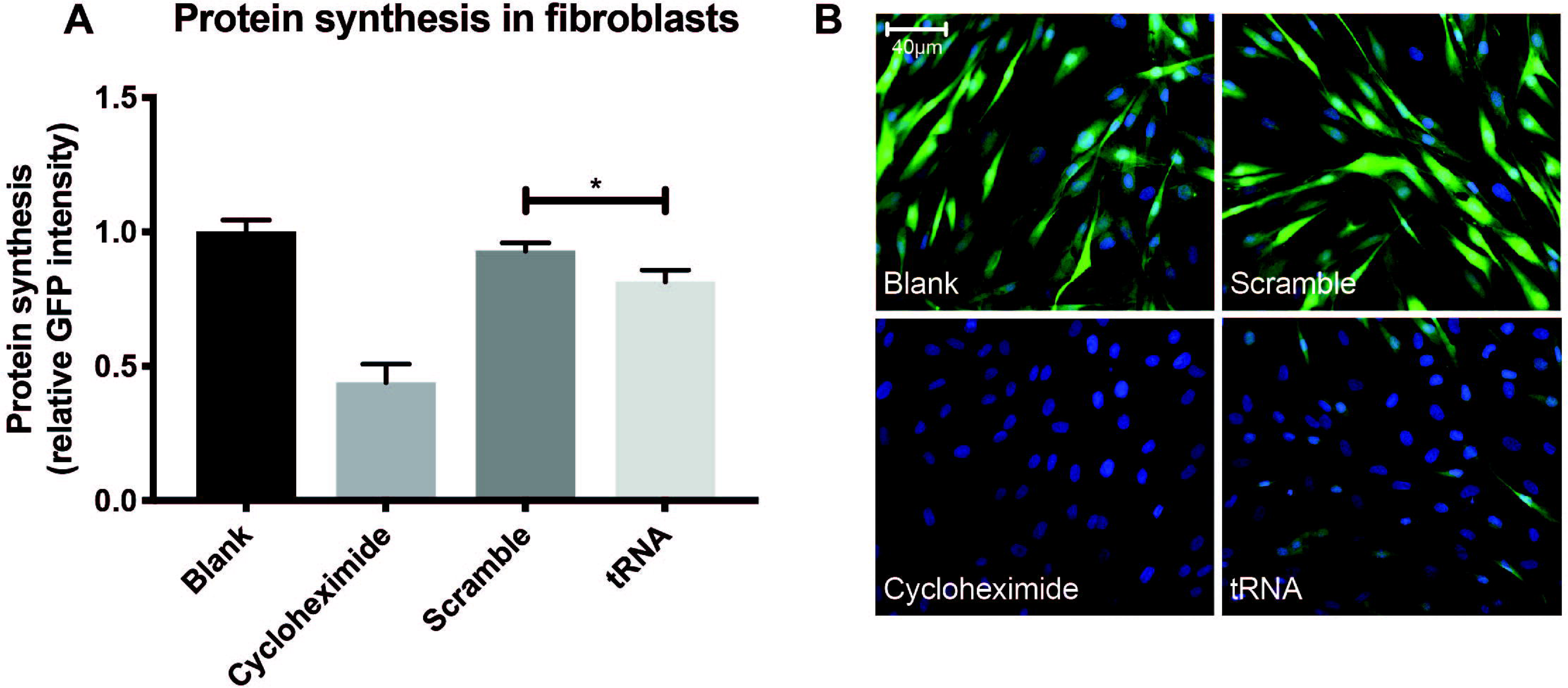
A: Green fluorescent protein (GFP) intensity relative to a blank (untreated) well in fibroblasts grown in medium with a GFP-labelled methionine analogue to quantify global protein synthesis. Cells were treated for 22 hours with cycloheximide (40 micrograms/ml), 500 nM tRNA-Gly-GCC and 500 nM tRNA-scramble. Data shown are mean ± SEM for 9 technical and two biological repeats. * = p <0.05. B: Representative images obtained using fluorescence microscopy and stained for newly synthesised protein (green) and DNA (blue) demonstrating no effect of 500nM tRNA-scramble on protein synthesis, but suppression of protein synthesis with cycloheximide (40 micrograms/ml) and 500nM tRNA-Gly-GCC.

## Discussion

This is the first study of tRNA species in syncytiotrophoblast-derived extracellular vesicles. We show they are the predominant form (>95%) of short RNA within STB-EV, compared to <50% within whole placental tissue, suggesting that tRNA is actively exported from the placenta via STB-EV. tRNA species were differentially expressed in SEV and MLEV, implying that the two vesicle types may play different roles in feto-maternal communication [1]. Most tRNA species identified within STB-EV were 5’-tRNA halves. The relative paucity of 3’-tRNA halves suggests the export of 5’-tRNA halves may be targeted for distinct biological functions.

5’-tRNA halves have previously been reported to inhibit both transcription (acting as miRNAs) and also translation [9]. Consistent with previous observations we show suppression of global protein synthesis in human fibroblasts by addition of exogenous 5’-tRNA-Gly-GCC. Normal human pregnancy requires the coordination of a myriad of physiological adaptations; STB-EV have been reported to be detected from as early as 6 weeks gestation and increase with gestational age [2]. Taken together, these data might suggest a novel role for 5’-tRNA halves within STB-EV in feto-maternal signalling in normal pregnancy.

5’-tRNA halves have recently been reported in extracellular vesicles derived from other human tissues, e.g. human seminal exosomes, which contain tRNA that are enriched for 5’ ends of 30-34 nt in length [12]. Murine epididymosomes differently express tRNA species in proximal and distal ends of the epididymis; the more abundant specific 5’-tRNA halves are similar to those we report in STB-EV (e.g. 5’-tRNA-half-GlyGCC) [13]. Others have identified tRNA halves in murine serum (bound to protein complexes), T cell fragments, yeast and *Drosophila* [14,15,26,27].

Translational arrest in cells where tRNA halves are produced is documented and involves displacing eiF4E from the translation initiation complex at the ribosome [10,16]. tRNA halves associate with argonaute proteins and inhibit transcription by acting as miRNAs which have seed sequences complementary to RNA targets [17]. 5’-tRNA halves can suppress around 70 genes relating to an endogenous retro-element target (MERVL) in cultured embryonic stem cells [13]. Given the variety of tRNA species identified within STBEV, we elected to investigate the *in vitro* effects of one of the most abundant (5’-tRNA-half-GlyGCC). Consistent with previous observations we demonstrated that a single 5’-tRNA half can have a profound effect on global cellular function in fibroblasts *in vitro.*

Different STB-EV types have distinct modes of cellular production and are postulated to have separate functions. We have identified not only a diverse array of tRNA species within STB-EV, but also different patterns of tRNA expression between different vesicle types (MLEV *versus* SEV), supporting the hypothesis that they may have different roles. STB-EV, in which 5’-tRNA halves are packaged, can be targeted to specific cell types, and to specific subcellular regions, by surface protein-protein interactions [6]. The variety of tRNA species we identified within STB-EV could thus potentially regulate transcription and translation more specifically at both a tissue as well as subcellular level. STB-EV are implicated in physiological changes seen in normal pregnancy, and drive pathological changes seen in diseases such as gestational diabetes and preeclampsia, where vesicle number and subtype are altered [28,29]. tRNA species contained within STB-EV might contribute to these processes.

The strengths of this study lie in its use of multiple techniques to interrogate the RNA content of STB-EV, as well as the demonstration of functional and descriptive data. Moving forward, one challenge will be isolating the contributions of individual tRNA species in vesicles that contain abundant quantities of diverse tRNAs. Future work should look to determine whether differentially targeted vesicles have different tRNA content, and whether tRNA content is altered in placental stress, such as that seen in preeclampsia.

In summary, we characterised tRNA species within STB-EV in normal pregnancy and found differential enrichment of individual tRNA species between MLEV and SEV. Most of these are 5’-tRNA halves, which was confirmed by qPCR. There is commonality of the most abundant tRNA species between patient samples as well as heterogeneity in the less abundant tRNA species. Finally, using an *in vitro* model, we demonstrated that 5’-tRNA halves can interfere with protein synthesis, suggesting they may play a role in feto-maternal signalling.

## Acknowledgements

We are grateful to Professor Dennis Lo at the Chinese University of Hong Kong for his critical appraisal of the manuscript. We acknowledge midwives Linda Holden and Fenella Roseman for recruiting patients.

## Funding

This work was supported by the National Institute for Health Research (WC and AC) and Medical Sciences Internal Fund, University of Oxford [grant number 0005271].

